# Example Based Hebbian Learning may be sufficient to support Human Intelligence

**DOI:** 10.1101/758375

**Authors:** Eric C. Wong

## Abstract

In this hypothesis paper we argue that when driven by example behavior, a simple Hebbian learning mechanism can form the core of a computational theory of learning that can support both low level learning and the development of human level intelligence. We show that when driven by example behavior Hebbian learning rules can support semantic, episodic and procedural memory. For humans, we hypothesize that the abilities to manipulate an off-line world model and to abstract using language allow for the generation and communication of rich example behavior, and thereby support human learning and a gradual increase of collective human intelligence across generations. We also compare the properties of Example Based Hebbian (EBH) learning with those of backpropagation-based learning and argue that the EBH mechanism is more consistent with observed characteristics of human learning.

## Introduction

From a computational perspective, it is not yet clear what basic learning rules are implemented by the brain to learn behaviors and store information. Simple local (Hebbian (Hebb, 1949)) learning mechanisms such as Spike timing dependent plasticity (STDP) (Song and Abbott, 2001) are well established biologically, but have not yet been shown to support learning of advanced tasks such as playing chess. In machine learning, artificial neural networks have been able to demonstrate good performance using learning rules such as backpropagation (Rumelhart et al., 1986) that can propagate training signals through deep networks. This success has prompted a search for biological learning mechanisms that are analogous to backpropagation (Bengio et al., 2015;Lillicrap et al., 2016;Guerguiev et al., 2017;Detorakis et al., 2019;Lansdell et al., 2019;Richards and Lillicrap, 2019) but the existence of such mechanisms has not yet been established. We argue here that simple Hebbian learning mechanisms are sufficient to support most types of learning in the brain and are also consistent with the way humans learn advanced behaviors.

In many analyses of Hebbian learning, synaptic plasticity is analyzed in the context of a neuron with a set of inputs, as shown in **Figure 1a**. The inputs are typically described by some statistical properties, and after Hebbian learning the activity in the output neuron reflected some property of the inputs, such as the projection of a sample onto the principal component of the input data (Oja, 1982). In Hebbian learning as implemented by STDP, it is necessary for presynaptic activity to precede postsynaptic firing in order to cause an increase in synaptic weight at a particular synapse. However, this does not imply that the presynaptic activity must have had a causal relationship with the postsynaptic activity, only that it *could* have a causal relationship. We focus here on the potential utility of Hebbian plasticity in a setting where the postsynaptic activity is determined primarily by inputs other than those undergoing plasticity. This generic scenario is shown in **Figure 1b**.

**Figure 1:**
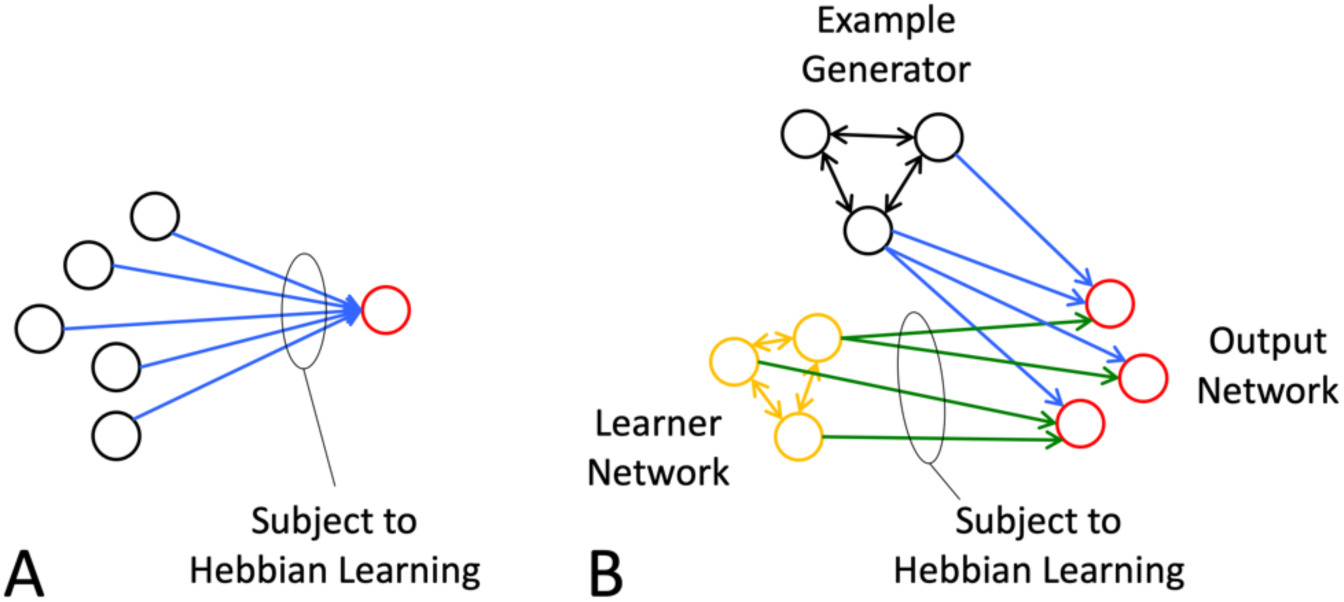
A) Common configuration for analysis of the Hebbian neuron. B) In Example Based Hebbian (EBH) learning, output is determined by a population of neurons here labeled the Example Generator. Hebbian learning in the synapses connecting a learner network to the same output neurons adjusts the synaptic weights such that the learner network can produce a similar output in the future.

The desired postsynaptic activity is determined by a source of example output, here referred to as the example generator (EG), which is assumed to be capable of producing the desired output through an output network (blue pathway in **Figure 1b**). In this model the system strengthens or creates by Hebbian learning new causal relationships between a learner network and the example output. The learner network produces some pattern of activity that may or may not initially be helpful in producing the desired output. During learning the EG produces the desired pattern of activity in the output network. If a Hebbian STDP type of learning rule is active at the synapses between the output of the learner network and the output neurons (green weights in **Figure 1b**), then those synapses whose pre-synaptic neurons became active just prior to the post-synaptic activity will be increased in weight. By this mechanism, the learner network will tend towards having a causal effect on the output network to generate the desired output in the future. We will refer here to this category of learning schemes as Example Based Hebbian (EBH) learning. Importantly, when an output neuron has two distinct and independent sets of inputs, as in this learning category, the behavior of the Hebbian neuron is qualitatively different. The learning that takes place is no longer well described by statistical properties of the total input stream, and will not reflect summary statistics, such as the principal components. Several learning strategies that would fall within the EBH learning category have been explored in previous work. The linear associator (Anderson, 1983) is a prototypical example of EBH learning, and this early work provides the underlying mathematical foundation reviewed in the Theory section below. Classical conditioning can be supported by Hebbian learning of the EBH type (Barto and Sutton, 1982;Tesauro, 1986) in which the conditioned stimulus is processed through the learner network, the unconditioned stimulus is provide by the EG, and the unconditioned response is the output. In fact, the observation of classical conditioning provides evidence that a mechanism that is functionally equivalent to EBH learning is active in the subjects of those experiments. In Contrastive Hebbian learning (Movellan, 1991;Xie and Seung, 2003;Detorakis et al., 2019), the output is held clamped while Hebbian learning is in effect throughout a network with both feedforward and feedback connections. This approach has been shown in simulation to be similar in performance to backpropagation under some circumstances (Movellan, 1991;Xie and Seung, 2003;Verstraeten et al., 2007;Detorakis et al., 2019), but requires feedback connections with uncertain biological plausibility. In reservoir computing (Buonomano and Merzenich, 1995;Verstraeten et al., 2007) a fixed network processes input and an output layer is trained using supervised learning, which may use Hebbian learning rules. In a recent example of reservoir computing of the EBH type, Illing (Illing et al., 2019) used a biologically plausible learning scheme in which the task was MNIST digit classification. Input images were sent to a hidden layer, with fixed random projections to a supervised output layer which was implemented as a spiking network, and subject to STDP plasticity. In this setting the Hebbian learning scheme was found to be comparable in performance to a non-convolutional network of similar size that was trained using backpropagation.

We point out in the following that EBH learning can provide a very general mechanism for attaching a desired behavior or memorandum to an arbitrary input. We then provide specific examples of EBH learning in support of episodic, semantic, and procedural memory, discuss the potential role of EBH in higher learning in humans, and compare the properties of EBH learning to those of backpropagation-based learning rules.

We hypothesize:

1. That a simple Example Based Hebbian learning rule for synaptic plasticity is sufficient to support both low- and high-level learning and is a core component of learning in the brain.
2. That the key to human intelligence lies in the development of an off-line world model which serves as a rich example generator for EBH learning.
3. That mechanisms for synaptic plasticity that propagate teaching signals through deep networks, as typified by backpropagation, are neither necessary nor likely in the human brain.

### Theory

We begin by noting that under simplifying assumptions EBH learning can provide exact learning of a desired association with a single training example. We consider a fully connected network of nodes with signed weights and linear activation functions, whose state at time *t* is represented by an activity vector *A*_*t*_. Let *A*_*t*_ be the input or the preprocessed input and assume that at time *t+1* the EG generates the desired output, described by the activity vector *A*_*t+1*_. Following (Anderson, 1983), a simple Hebbian learning rule produces a change in synaptic weights W at the time of the transition that is given by a learning rate *λ* times the outer product of the activity vectors:

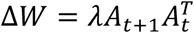

Assuming zero initial synaptic weights *W*, the weight matrix after this transition would be equal to *ΔW*, and the next time that the input *A*_*t*_ is introduced to the network, the network would produce in the next time step:

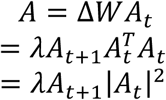

Note that both *λ* and |*A*_*t*_|^2^ are scalars, and the output is therefore an exact copy of *A*_*t+1*_ apart from scaling. If in addition the learning rate is set such that *λ*|*A*_*t*_|^2^ = 1, then the output *A*_*t+1*_ is reproduced exactly in response to the input *A*_*t*_. In practice, several factors will make this reproduction imperfect, including the fact that the weights learned in the transitions between different pairs of states will in general overlap and confound one another, that biological networks are generally connected sparsely rather than fully, and that the activation function is not linear. Nevertheless, this demonstrates the idea that simple Hebbian learning can approximately attach an arbitrary output to an arbitrary input and can do so from a single example with an appropriate learning rate.

We next provide three examples in which the EBH learning mechanism can provide learning in support of episodic, semantic, and procedural memory. These examples are simple in construction, and are intended to demonstrate the range of basic learning types that can in principle be supported by an EBH learning structure, rather than to represent detailed realistic models. More detailed but important considerations such as mechanisms for constructing attractors (Fiebig and Lansner, 2017), for maintaining criticality (Deco and Jirsa, 2012;Cocchi et al., 2017), and for promoting and maintaining sparsity (Rozell et al., 2008;Wadhwa and Madhow, 2016) are not considered here. The code for these examples is at: https://github.com/spindragon/EBH_Learning, and more details can be found in the code.

#### Episodic Memory

EBH learning can record a series of engrams as an ordered list

In the episodic memory system of mammals, it is thought that experiences are processed and encoded into patterns of activity in the hippocampus known as engrams (Josselyn, 2010;Tonegawa et al., 2018). These engrams arrive in the hippocampus as a stream and are apparently recorded on a single shot basis as each one would typically occur only once. The engrams are thought to both represent an event or state for the purpose of short term memory and also have the ability to evoke responses to the memory of the state (Liu et al., 2012). The EBH learning framework provides a simple mechanism for this stream to be recorded as a linked list, such that in the future, evoking one engram in the list naturally generates the stimulation of the subsequent engrams, in forward time order. This can be accomplished simply by employing an EBH mechanism continuously throughout a network that receives a series of engrams as externally specified patterns of activity. Similar networks were analyzed by Kleinfeld (Kleinfeld and Sompolinsky, 1988).

As an example, we simulated a binary network of 10,000 nodes, connected by sparse connections with a random connection density of 2%, initialized with zero weights. Into this network we introduced a series of 26 engrams, each consisting of approximately 150 active nodes, depicted in Figure 2 as 150 active pixels on a background of 10,000 pixels, with each engram arranged in the form of a letter on a 100 x 100 grid only for recognizability. At each transition between engrams,

**Figure 2:**
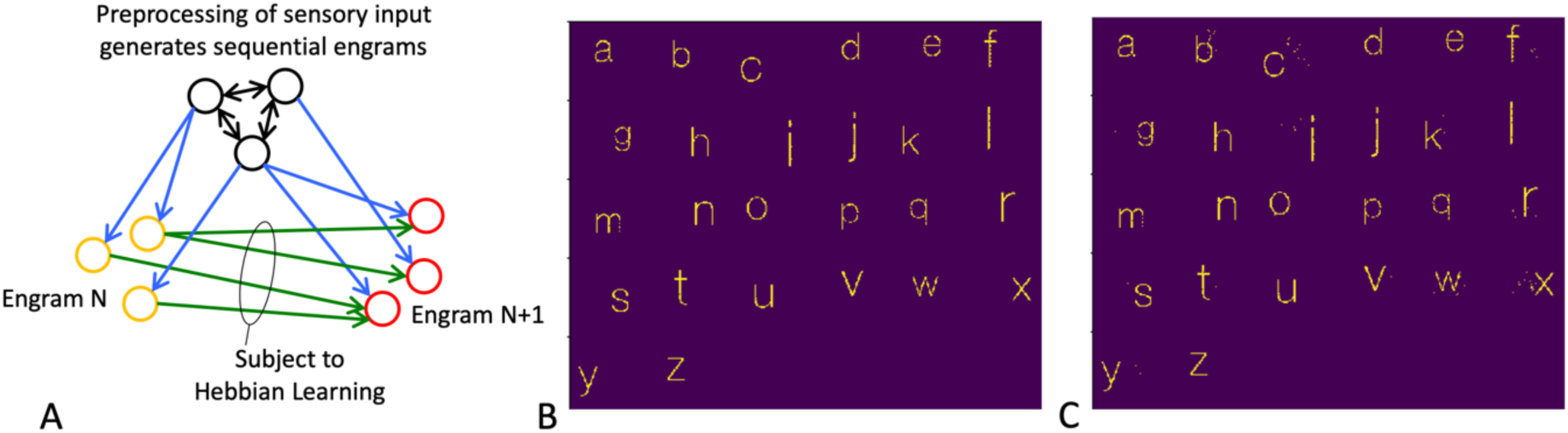
EBH supports single shot learning of episodic memory as a linked list. A) In this example, the EG is assumed to be a network that preprocesses input and cognition into a series of engrams to be recorded. B) Within each 100×100 patch of pixels, the pixels in yellow represent a sparse engram, represented as a letter only for recognizability. The engrams are introduced to a naïve network sequentially, with Hebbian learning continuously active at all synapses. C) After learning, the first engram was reintroduced to the network, and sequentially recreates the original series of engrams with some errors (shown as spurious pixels).

Hebbian learning was simulated by adjusting the weights in proportion to the outer product of the pair of engrams, as in the Theory section above. In this case the activation function was a step function in order to keep the activations binary, with a fixed threshold that was set to produce approximately 150 active nodes at each step. After a single learning pass of the 26 engrams, the network was tested by reintroducing the first engram (‘a’) and running the network through 25 forward time steps, resulting in an approximate reproduction of the series of engrams in forward time order (**Figure 2**). Initializing the network with any of the learned engrams resulted in the network running through the remaining engrams in order. This example demonstrates the use of Hebbian learning to enable single shot recording of a sequence of neuronal states, resulting in a recallable ordered list. The strong directionality in the recollection of the sequence is consistent with our experience that sequences are much easier to recall forward than backward.

#### Semantic Memory

Learning multiple attributes of an object

Semantic memory describes another broad category of knowledge that can be learned, and refers to knowledge of objects, concepts, and their relationships. As an example of learning in this category, we describe here extensions of the episodic memory example above to demonstrate learning of multiple attributes of a set of objects. For the set of English letters, we demonstrate learning of the next letter in the alphabet, as above, but also the letter of the opposite case, as well as learning which case each letter is an example of (**Figure 3**). As above, each letter is coded as a set of binary nodes in a network of 10,000 nodes, arranged for convenience as the letter itself when displayed on a 100×100 grid. In addition, a small set of nodes (50) was activated to instruct the network to provide the next letter, the letter of the opposite case, or an indication of the case of the current letter. These instruction and output nodes are shown in **Figure 3B**. Each of the three tasks (next letter, change case, and identify case), was trained once, using as input one of the instruction codes in addition to a letter, and with the correct output as the teaching signal of the EG (**Figure 3A**). For testing, the letters and instructions were input to the network, and the output of the network was determined using a winner take all (WTA) approach in which the response was taken to be the output with the largest inner product with the observed network activity. Results are shown in **Figure 3C-D**. This example demonstrates that the EBH mechanism can learn multiple attributes of a single object (next letter and opposite case) as well as a many-to-few mapping, as in the identification of the case.

**Figure 3:**
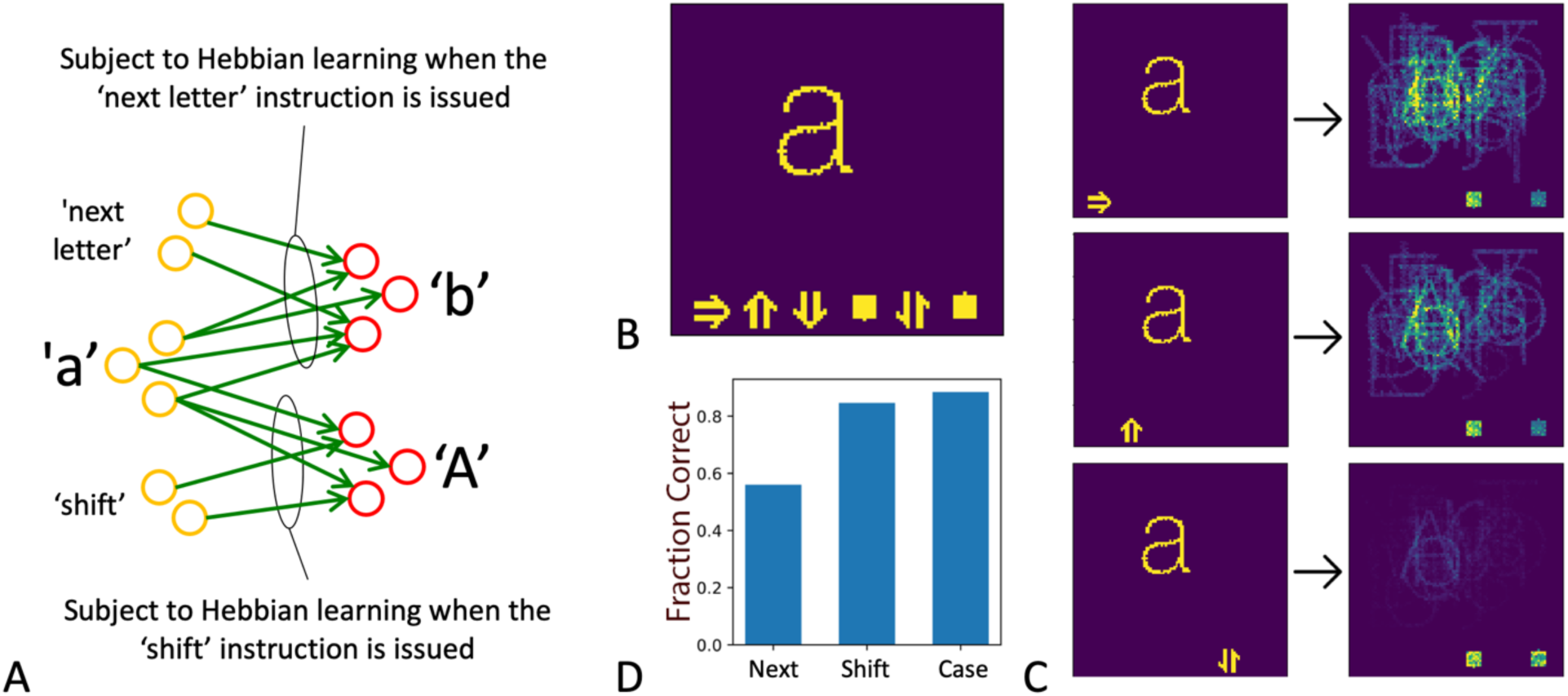
EBH learning of multiple associations (see text). A) For each input letter (here ‘a’) the network learns multiple associations. For each combination of input letter and instruction, the EG (not shown in figure) is assumed to produce the correct output. B) In addition to the input letter, six additional sets of 50 nodes each are defined and shown along the bottom arranged as symbols. These are used to provide instructions to the network and responses to queries. From left the symbols are designated: next letter; shift, convert to lower case, lower case response, query case, and upper case response. C) When ‘a’ and the ‘next letter’ instruction are applied to the trained network, the response from the network is shown, and the inner product of the response with all letters shows that ‘b’ is prominent within the response. In this case ‘b’ has the highest correlation with the response among all possible letters and is therefore the WTA response of the network. With a shift instruction, ‘A’ becomes prominent in the response, and with a case query, the lower case response is slightly more prominent than the upper case response. D) Correct response rates for all three tasks. With random responses the correct response rate would be 1/52, or approximately 0.02 for ‘next letter’ and ‘shift’ tasks, and 0.5 for the ‘case query’ task.

#### Procedural Memory

EBH learning can attach a sequence of actions to a reproducible source of reference activity

In reservoir computing, a network (the reservoir) provides a pattern of network activity, and the desired behavior is attached to that activity. The reservoir may provide preprocessing of input data (Buonomano and Merzenich, 1995;Verstraeten et al., 2007) or provide a time base for coordination of movement during dynamic motor activity. In the hippocampus there are thought to be cells or networks that are responsible for producing a time reference (MacDonald et al., 2011;Eichenbaum, 2014;Salz et al., 2016), and in this example we demonstrate that when a reproducible source of time synchronized neuronal activity is available, EBH can attach a sequence of motor outputs to the time signal. The structure of the network in this example is a random but fixed reservoir network acting as a time base, with Hebbian learning at the output (**Figure 4**). The output is initially controlled by an EG such as the conscious execution of a sequence of motor movements, and the goal is to learn the movement so that it can be automatically generated in the future (**Figure 4A**). A humanoid figure was simulated in 2D space with human-like dimensions and weights, but with only a torso and legs. The physics of movement was simulated by the Pymunk physics engine, and the figure was controlled through 6 torques, applied at the left and right hip, knee, and ankle joints. A sequence of torques that resulted in the execution of a backflip was empirically hand coded and used as the EG (**Figure 4B**). The movement was executed over 1.25 seconds with 10ms time resolution. With the reservoir initialized to a known initial state and fixed random weights, the sequence of output torques was executed while EBH learning was active in the weights connecting the reservoir with the output. After a single example of a backflip by the EG, the movement is approximately contained in the output weights, with errors arising from synapses that may be used for multiple steps in the motor sequence. After learning, the reservoir was reset to the same initial state and allowed to run, with the motor output being driven by the reservoir and the learned output weights. The resulting movement is shown in **Figure 4C**, and closely approximates the EG. While biological motor planning, coordination, execution, and correction are known to be much more complex (Svoboda and Li, 2018), the example shown here is intended to demonstrate the approximate attachment of a desired sequence of behaviors to an underlying fixed, repeatable sequence of activity from another network.

**Figure 4:**
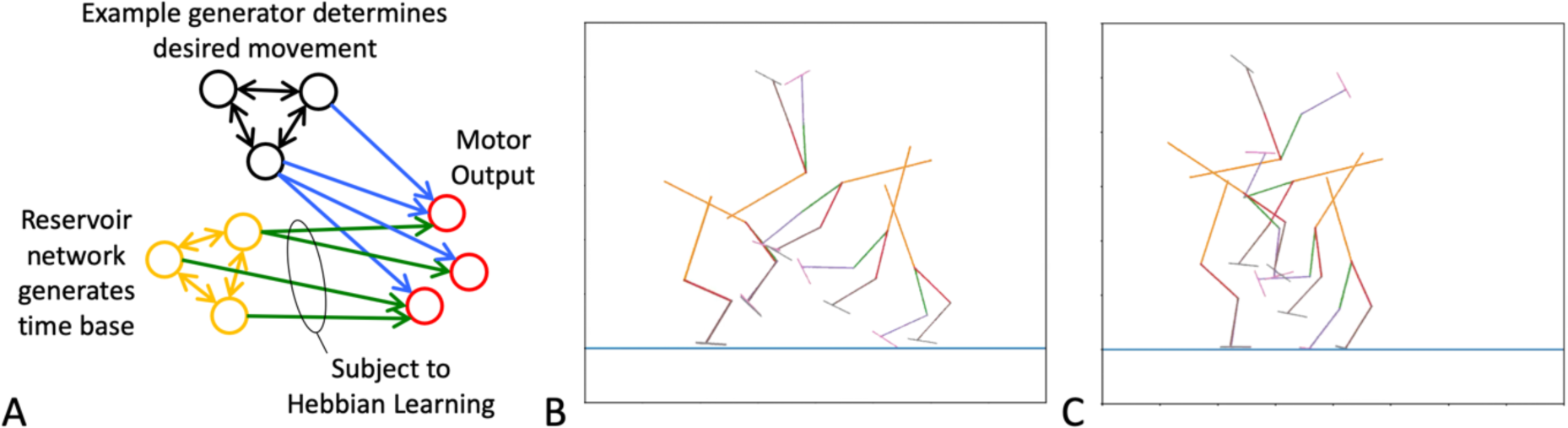
EBH learning of a dynamic pattern of activity. A) In this example conscious motor control is the EG and produces example motor output. A reservoir network serves as the learner network and generates a reproducible time base. B) a 2D simulated anthropomorphic figure was empirically programmed to perform a backflip. Six frames from the maneuver are shown. C) After one training example, the reservoir generates the same time base, and the output determined by the newly learned weights produces a similar but not identical maneuver.

In addition to networks that are thought to provide a time base, there are also cells known as place cells, that together provide a spatial basis that appears to be stable (Muller, 1996;Smith and Mizumori, 2006;Moser et al., 2015). The EBH mechanism may also be used to attach information from landmarks in an environment to their locations as a new space is explored and mapped.

### From Prediction to Abstract Associations

The prediction of future events is thought to be a central function of the brain as it can improve decision making and enable anticipatory behavior (Schultz et al., 1997;Rao and Ballard, 1999;Friston, 2010). We note that in very simple configurations EBH learning directly converts evidence from the environment into predictions. Imagine a pair of neurons A and B in a primitive organism that respond to two different stimuli S_A_ and S_B_. If S_A_ consistently precedes S_B_ and there are silent synapses from A to B, EBH learning will strengthen those synapses, and in the future S_A_ will excite B, forming an internal connection by which S_A_ predicts S_B_. Classical conditioning also follows this same pattern, as the conditioned stimulus becomes a predictor of the unconditioned stimulus. We propose that EBH learning provides a mechanism for direct online conversion of evidence from the environment into predictions.

In lower animals, the data that is converted into predictions comes directly from the environment, and the predictions themselves form an implicit model of how the world works. While this assures that learned predictions are based on evidence from the physical world, it also limits associations to those that have been directly experienced. In humans, learning has progressed to the construction of explicit models of the world that can be manipulated off-line (Lake et al., 2017). This off-line world model, also known as imagination (Asma, 2017;Vyshedskiy, 2019) and here defined as the ability to think about objects and events not tied to the physical present, allows for hypothetical events and actions to be considered. In addition, the development of language has allowed for objects, actions, and ideas to be abstracted into words and communicated from person to person (Gleitman and Papafragou, 2005). We hypothesize that with the benefit of language and an off-line world model, EBH learning can support much of human intelligence by enabling the association of objects and actions through the use of abstract, hypothetical, and complex example behavior. For humans, the EG can include instruction from others, detailed mimicry of others, and ideas from conscious thought, in addition to direct evidence from the environment. We propose that the core physiological learning mechanism (EBH learning) may be the same in humans as it is in lower animals, but that the richness of potential sources for the EG is a key distinguishing feature of human intelligence.

As a simple example, a child may learn that the word ‘three’ is associated with the symbol ‘3’. A teacher would present an image of the numeral ‘3’ and say the word ‘three’ to the child. If the child has learned to mimic, she would repeat the word ‘three’ along with the teacher, and as in classical conditioning the word would become associated with the numeral. Later in school, the child might learn the Pythagorean Theorem through the same process. The formula a^2^+b^2^=c^2^ would be attached to the concept of a right triangle with sides and hypotenuse a, b, and c. Both the conditions for the theorem and the theorem itself are significant abstractions relative to what most animals are capable of, but when learning this theorem in school, this attachment would be a straightforward application of EBH learning and represents one step in the incremental learning of associations along the mathematical curriculum.

We hypothesize that the EBH learning system initially evolved to directly convert evidence from the physical environment into learned predictions, and subsequently adapted to produce arbitrary associations, mostly communicated by other humans. This evolution from prediction as an implicit world model to arbitrary associations within an off-line model would have come with two major competitive advantages, and one major disadvantage. First, the ability to consider hypothetical scenarios and predict hypothetical consequences enables longer term planning than is possible without this facility and may have enabled the human path to successful adaptation to changes in environmental factors such as climate or ecosystems. Second, the EBH learning mechanism in conjunction with language provides a simple and direct way to transfer associations from person to person, thereby enabling accumulation of human expertise and knowledge across generations. Indeed, much of the early life of modern humans is spent setting in place a large library of associations that sample the collective knowledge of human history through both formal and informal education. A major disadvantage of enabling the learning of arbitrary associations is that the associations are no longer necessarily tied to evidence from the environment, thus allowing for both facts and misinformation to propagate.

### EBH vs Backpropagation

In recent years machine learning has seen tremendous success in areas such as image classification (Krizhevsky et al., 2012;He et al., 2015) and game play (Mnih et al., 2015;Silver et al., 2018) using deep networks with credit assignment calculated by backpropagation. As these networks are loosely based on the architecture of the brain, it is natural to ask whether the brain may also perform credit assignment using backpropagation or some method that is functionally similar to backpropagation, and several models within this class have been proposed (Bengio et al., 2015;Lillicrap et al., 2016;Guerguiev et al., 2017;Detorakis et al., 2019;Lansdell et al., 2019;Richards and Lillicrap, 2019). In backpropagation, an error signal (or reward) is determined at the output of a network, and with knowledge of the inputs, synaptic weights, and activation functions of the network, the dependence of the error on each of the synaptic weights can be calculated and used to adjust all weights so as to reduce the error. Here we contrast EBH learning not with a specific backpropagation-based algorithm, but with the general class of learning methods that uses end-to-end adjustment of synaptic weights to improve network performance, of which backpropagation is a prototype, and we use the term backpropagation to describe this class for convenience. We outline here two major ways in which EBH learning and backpropagation are different, and the potential impact of these differences on the likely role of each in the human brain.

First, we contrast the basic premise of each learning mechanism. For EBH learning, expertise is gained in incremental steps, with some source of explicit example behavior, but the learning of each step may occur in a single shot. The learning mechanism itself is very unsophisticated and simply records associations or behaviors that are identified by the EG and makes them available in the future. The intelligence of the resultant network is derived primarily from the EGs, whether from teaching by others, observation of others, or results from internal thought. With backpropagation, a cost function is defined and the backpropagation algorithm gradually modifies the synaptic weights to improve performance. Intelligence is actually generated by the backpropagation algorithm itself. The learning mechanism ‘figured out’ how to solve the problem, but because the intelligence was generated by the set of rules that guide the synaptic plasticity, it is not clear how the organism would gain any understanding of how the problem was solved. We argue that EBH learning is generally a better fit to the way humans learn than backpropagation. In the example above of learning the Pythagorean theorem, the acquisition of expertise in using the theorem is an abstract but simple association that was provided by an instructor, it is easy to imagine that it could be committed to memory in a single example, and the theorem is one of many small, incremental steps along a math curriculum, all traits that are consistent with EBH learning. The incremental and explicit steps along the curriculum provide identifiable units of information and represent the understanding of the subject in the sense that they can be passed from person to person. As a different example, take a game such as chess or go, in which machine learning methods that employ backpropagation have been extremely successful. With Alpha Zero (Silver et al., 2018), the machine learning algorithm progressed from naive to super-human levels in both chess and go in a few days with only self-play, given no domain specific knowledge other than the rules of the game. As examples of advanced human tasks that were mastered using backpropagation-based methods, these could be seen as examples that might support the possibility of biological backpropagation. However, expert human players of these games learn largely from extensive study of the games and analysis of previous experts, and can describe strategies, teach students using an incremental curriculum, and provide analysis. It is difficult to imagine how a network that is globally optimized for a complex task using a backpropagation type approach could articulate what principles and strategies were learned, and how to incrementally teach another human this expertise.

We argue that the vast majority of what each contemporary human knows is learned from the existing body of human knowledge, and that the accumulation and communication of knowledge and expertise across generations is a hallmark of human intelligence. However, it is also true that individuals can innovate and produce expertise that they are not given. The backpropagation mechanism can innovate, and it is in fact incumbent upon the backpropagation algorithm to discover solutions to the optimization problems it is given. However, it is not clear, as noted above, that new expertise gained through backpropagation could be described in a way that can be communicated and contributed to the body of human knowledge, unless the innovations themselves are understood by study after the fact. New expertise gained by backpropagation would by definition be experienced as intuition, as it is not gained through conscious reasoning. For an organism that uses EBH learning, the process of innovation would be quite different. The EBH mechanism itself does not innovate, but simply sets associations defined by the EG in place in the network. Under the EBH hypothesis two types of innovation could occur. One is that the simple act of setting in place many associations within the same network would naturally create by chance many input/output mappings in addition to those specifically trained by the EG. These mappings would be likely to effectively implement algorithms that summarize those associations, and potentially also allow for some level of generalization and analogy. This type of innovation would also be experienced as intuition, since it is also not derived from conscious reasoning. A second type of innovation within the EBH framework that may arise from within an organism is revelations from conscious thought. Our ability to consider hypothetical situations, ideas, and questions in an off-line model of the world allows for such revelations to not only serve as EGs for the individual, but they can also typically be articulated and communicated so that they can serve as external EGs for others and contribute to the collective human intelligence.

The overall picture we propose is that EBH learning can support several lower level learning mechanisms that support learning of intuitive knowledge across species, and that the same mechanism also serves to record knowledge and innovations that come from external examples and conscious reasoning in humans and perhaps some higher animals. Backpropagation is a very efficient means of generating new expertise but can only generate intuitive expertise.

A second major difference between EBH learning and backpropagation is that EBH learning is minimally invasive to the network as it establishes associations, while the backpropagation algorithm can restructure the entire network as it searches for solutions. In the limit of a densely connected network with all nodes active, both EBH learning and backpropagation will generally change all synapses. However, the activity in the brain is sparse (Shoham et al., 2006;Barth and Poulet, 2012). In this regime, the synaptic changes made by EBH learning, which is given by the outer product of the EG activity and the preceding network activity, both of which are sparse, would be very sparse. For example, if the typical number of active neurons in a network is a fraction S of the total neurons, the outer product of two activity vectors will have a fraction S^2^ of non-zero values. Thus if S is on the order of 0.01, the EBH learning mechanism will modify on the order of S^2^=0.0001 of the synapses in a network for each learning event. For backpropagation, this fraction is more difficult to estimate, as it depends in detail on the network structure and connectivity in addition to the activity sparsity S, but because it propagates through the layers of the network, each learning event will modify many more synaptic weights than EBH. The EBH learning mechanism is in effect shallow in the sense that it only affects one layer of the network, that layer which is located at the confluence of the signal paths of the EG and the learner network. However, in a network that is strongly recurrent this confluence is not necessarily at a fixed set of output nodes, and the EBH learning mechanism may in fact have access to most or all of the synapses throughout the network, though only a small fraction (S^2^) could be modified by a single learning event. In the episodic memory example above for example, the engrams, and therefore the modified synapses, were sparsely and randomly distributed throughout the network.

In the context of a large network that is trying to gain expertise in a vast number of different areas, this minimally invasive approach to learning each incremental component may be less likely to incur confusion between tasks. Backpropagation, by contrast, has more power to shape a network toward a particular goal, but at a greater risk of unlearning expertise that was previously acquired.

Regardless of whether an EBH learning mechanism, a backpropagation type mechanism, or some other mechanism is responsible for the coordination of synaptic plasticity, there are also well established neuromodulatory mechanisms in place to accept or reject candidate synaptic changes (see (Kusmierz et al., 2017) for a recent review). While these mechanisms provide an important reward-based feedback component to the learning system, it is not obvious that their presence helps to distinguish between EBH and backpropagation hypotheses.

## Discussion

We are interested here in identifying the core learning rule or rules that can serve as a foundation for understanding what guides synaptic plasticity and for building more complete and detailed computational models of human learning. We hypothesize that simple EBH learning mechanisms along with reinforcement from neuromodulatory feedback may be sufficient to provide this foundation. We argue that the key innovations that underlie human intelligence are the development of language and imagination in combination with simple EBH learning, and that together these allowed for the gradual increase of collective human intelligence. We point out that while backpropagation based rules can independently support advanced learning, there is no known mechanism by which expertise gained using backpropagation can be articulated in the way that humans communicate ideas.

The examples provided here are very simple and intended to point out broad classes of learning settings in which EBH may be useful. We have not shown here that EBH learning is capable of learning complex tasks, and this is an obvious next step. Candidate tasks for more complex computational simulations with EBH learning could be games such as chess where a large body of training data is available, and also data is available from backpropagation based approaches for comparison of features such as learning rates.

Another useful computational comparison between EBH learning and backpropagation based approaches will be the development of computational learning models that perform many tasks within one network. Within such networks, the dependence on curriculum (Bengio et al., 2009) the emergence of modularity and compositionality (Yang et al., 2019), and the emergence of analogical reasoning (Gentner and Holyoak, 1997;Krawczyk, 2012;Hobeika et al., 2016) may demonstrate whether these approaches have behavioral characteristics that parallel those of the brain.

Biological validation of the EBH learning hypothesis is also possible. For example, the EBH mechanism would predict that during learning, both the EG and the learner network are active, while after learning only the learner network is active. This is consistent with the experimental observation that the amount and distribution of activation in some areas of the brain decreases with increasing expertise (Flament et al., 1996;Toni et al., 1998;Halsband and Lange, 2006), but this observation is also consistent with any learning mechanism in which high activity levels are present during learning, and physiological mechanisms operate to increase sparsity and network efficiency over time after learning. The observation of STDP itself shows that EBH learning *can* happen, but not that it does. It is specifically the provision of a source of example behavior that distinguishes EBH learning. The demonstration of classical conditioning in neural preparations further demonstrates that a mechanism that is functionally equivalent to EBH actually *does* happen because an example (the unconditioned response) behavior is provided, and results in a conditioned response. In general, validation of the EBH model would involve demonstration that prior to learning, a source of neural activity that is identifiable as an EG produces a particular behavior, and that after learning a different source of neural activity produces the same behavior.

In this paper we outline what may be a core component of the learning mechanism for both higher and lower animals. For some low-level learning mechanisms such as the episodic memory example described above, the EBH mechanism may be sufficient by itself to perform the required learning. However, we have postulated that for human level learning, EBH is aided by a conscious control process which is critically dependent upon language for abstraction and communication, and upon the utilization of an off-line world model to enable hypothetical thinking and rapid generation of examples. We note that the evolutionary step of developing an imagination, or off-line world model, would have been a difficult one, as it is critical for survival to maintain a very clear real-time distinction between reality and imagination. The fact that schizophrenia, which is characterized by an inability to maintain this distinction, remains prevalent in modern humans (approximately 0.5% (Saha et al., 2005)) suggests that this distinction is indeed difficult to maintain reliably. The difficulty of this evolutionary step may be one reason that higher intelligence is not more widespread in the animal kingdom. The process by which this step took place is unclear, though we note that the process of developing an imagination does occur in all human infants around the age of 2 (Ganea et al., 2007), and study of this developmental step may provide insight to inform models of imaginary thought.

An important question for the EBH hypothesis is: what is the structure of this conscious control process, and how can it be described in a way that is biologically plausible and useful as a component of a computational model? We do not speculate on the answer to this question here, but note that this process is closely related to phenomena studied under many labels, including conscious control (Schneider and Shiffrin, 1977;Shepherd, 2015), imagination (Ayman-Nolley, 1992;Asma, 2017), working memory (Fiebig and Lansner, 2017;Velichkovsky, 2017;Masse et al., 2019), and consciousness (Dehaene and Naccache, 2001;Tallon-Baudry, 2011;Koch et al., 2016;Tononi et al., 2016;Velichkovsky, 2017), and prior work in these areas may provide clues as to how this process emerges and operates, and how it may contribute mechanistically to EBH learning.

## Acknowledgments

We would like to thank Richard Buxton, Thomas Liu, and Peter Bandettini for many valuable discussions regarding the subject of this work and for helpful review of this manuscript, Alan Simmons for helpful review of this manuscript, and Maxwell Wong for help in coding of the examples.

